# The effect of hydrophobic gases on the nervous system of Daphnia magna

**DOI:** 10.1101/2021.02.19.431892

**Authors:** K. Carlo Martín Robledo-Sánchez, J. C. Ruiz-Suárez

## Abstract

Hundreds of hydrophobic substances: alkanes, alcohols, benzodiacepines, barbiturates, ethers and even gases, can induce General Anaesthesia (GA) in mammal animals. Moreover, it has been suggested that the primary site of action of such agents is on the spinal cord. Here, we investigate a scenario that is in double measure important to understand the mechanisms behind GA: its induction under water in invertebrate animals. We evaluate the capacity of xenon, nitrous oxide and krypton to suppress harmful sensations, provoked by intense light, in the crustacean D. magna. Due to the hydrophobic characteristics of those gases, we designed a special chamber to force them to dissolve in water at pressures up to to 50 atmospheres, whereas at the same time measure in real time the motility of the animals. Surprisingly, the aquatic animals are immobilized with xenon and nitrous oxide. Under this condition, they don’t respond to a noxious stimulus. Our results are crucial to understand the action of inert gases in GA and the role of the spinal cord.

## 1 Introduction

An ideal anesthetic must confer a rapid and adequate anesthesia induction, effective depression of the autonomic nervous system, and analgesia without untoward effects. The most used anesthetic gas in clinical practice nowadays is isoflurane (C_3_H_2_ClF_5_O). This molecule possesses hydrophobic characteristics and a blood/gas partition coefficient of 1.4. Despite it is one of the most used anesthetics, its implementation in hospitals is cautionary due to multiply undesirable effects: hypotension, respiratory suppress, nausea, among many others. It has been claimed recently that the noble gas xenon (Xe) is much safer with no secondary effects. This noble gas is an element characterized by an outer shell filled with electrons, *i.e*., it possesses a null chemical reactivity. Xenon is a naturally occurring trace gaseous element with a concentration in the Earth’s atmosphere of 1 part per 11 million.^1^ Contrary to inhalation agents like enflurane, desflurane, and sevoflurane, that have an important impact on atmospheric pollution,^2–5^ the natural presence of xenon does not produce ozone-depletion and greenhouse effects. Also, it represents a considerable reduction of workplace risk and environmental hazards.

Xenon is an attractive element for pharmacology due the protective effect in several organs,^6–8^ with a greater regional perfusion and circulatory stability.^9,10^ Furthermore, this element poses low chemical reactivity with drugs^11^ and a blood/gas partition coefficient of 0.15.^12^ For this reason, xenon presents a rapid induction and a short recovery time, without negative effects.^13^

Xenon starts to be more used in clinical practice than nitrous oxide^14–16^, due to the comparative lower risk and its pharmacological profile, which is very similar for both gases (inhibition of N-methyl-D-aspartate). Its use, however, is limited due to the fact of the extended technical requirements it needs and to its high cost. Krypton also presents anesthetic effects but not under normal pressure conditions.^17^ This property has been studied in several animal models, although in several cases no unequivocal signs of narcosis could be demonstrated. The capacity of krypton to induce anesthesia has been poorly explored and non-clear results have been reported in normobaric conditions.

Nitrous oxide N2O (also known as laughing gas) is a gaseous compound with anesthetic and analgesic properties.^18^ The mechanism of its action consists in the inhibition of spinal cord dorsal horn neurons and does not present significative differences with xenon.^19^ Nevertheless, nitrous oxide in contrast with xenon can be a health hazard compound after long time exposure.^20^

The crustacean *Daphnia magna* (DM) has been previously defined as a potential model to test the potency of inhaled anesthetics (halothane, isoflurane and enflurane).^21^ Furthermore, DM have been used to test the immobilizer effects of local anesthetics.^22^ The invertebrate DM’s nervous system shares several similitudes with mammals, as the presence of several neurotransmitters like acetylcholine, serotonin, dopamine, epinephrine and GABA receptor signaling pathways.^23^ The aim of this study is to carry out a series of experiments to test the potency of three gases to induce a state of anesthesia in an invertebrate model. We consider this aim of great importance because our findings could discern if the primary target to induce immobilization in animals is in the brain stem or spinal cord.

DM was exposed to hyperbaric conditions of xenon, nitrous oxide, and krypton. The pressures used were 1-14 bars for xenon, 1-16 bars for nitrous oxide and 1-44 for krypton. The motor response of the daphinds after a harmful stimulus produced by an intense red light was evaluated by calculating the autocorrelation of swimming images, which corresponds to changes in the position of all organisms in each frame respect to the original positions. By using this method, we are able to take into account the complete animal population, see the movie in the Supplementary Material.

## 2 Materials and methods

### 2.1 Ethics

The committee of animal care and use of the Center for Research and Advanced Studies approved the protocol, and the study was performed in accordance with relevant aspects of ARRIVE guidelines. No specific permissions were required for experimentation.

### 2.2 Daphnia magna

DMs were acquired from a permanent pond in Guadalajara, Mexico. The DMs were acclimated and cultured in the laboratory during 3 weeks at 21 ± 2 °C under a 12-h light: 12-h dark photoperiod and fed daily with unicellular culture *Chlorella vulgaris*.^24^ The cultured consist of a 1-liter glass beakers with 800 ml of a particular physiochemical characteristics medium (reconstituted water). This liquid medium was prepared with: 25 mM NaCl, 2 mM KCL, 1 mM MgCl_2_, 2 mM CaCl_2_, 2 mM MOPS as a pH buffer, and 1 mM NaHCO_3_.^25^ Medium pH and hardness CaCO_3_, was monitoring once a week (Thermoscientific Model Versastar Pro, USA). Dissolved Oxygen (DO) was determined in agreement to Mexican Official Norm, after 48 hours of aeration. Reconstituted water was refreshed twice a week and when necessary we replenished the lost volume by evaporation with miliQ water.

### 2.3 Gases

Three gases were used to induce anesthesia on DM: xenon (investigation grade 5.5, CAS-No. 7440-63-3, analytical standard, purity >99%), krypton (investigation grade 5, CAS-No. 7439-90-9, analytical standard, purity >99%), nitrous oxide (CAS-No. 10024-97-2, clinical standard, purity >99.5%), which were acquired from Praxair Technology Inc (Mexico). All others chemical used were analytical grade and were purchased from Sigma-Aldrich (USA). The criteria selection for experimentation with these gases were: (1) low chemical reactivity, (2) slight to no hazard of toxicity and untoward effects, (3) high coefficient partition oil/water, (4) low coefficient partition blood/gas. To evaluate the pressure effect on the animals, nitrogen gas (CAS-No. 7440-63-3, analytical standard, purity >95%) was used as a control.

### 2.4 Pressure tolerance assay

DM was subjected to hyperbaric conditions of nitrogen to evaluate survival. For this experiment, six daphnids 2-4 weeks old (2.5-3.4 mm in length) were exposed to different pressures for 1 hour. Daphnids were introduced in an open micro centrifuge tube filled with 1.5 ml of reconstituted water. The micro centrifuge tube was placed in a cylindrical device and hermetically sealed, see Figure 1a. The criteria to discern if the DM survive to high pressures was to consider their mobility, cardiac contractions and tissue damage.

**Figure 1.**
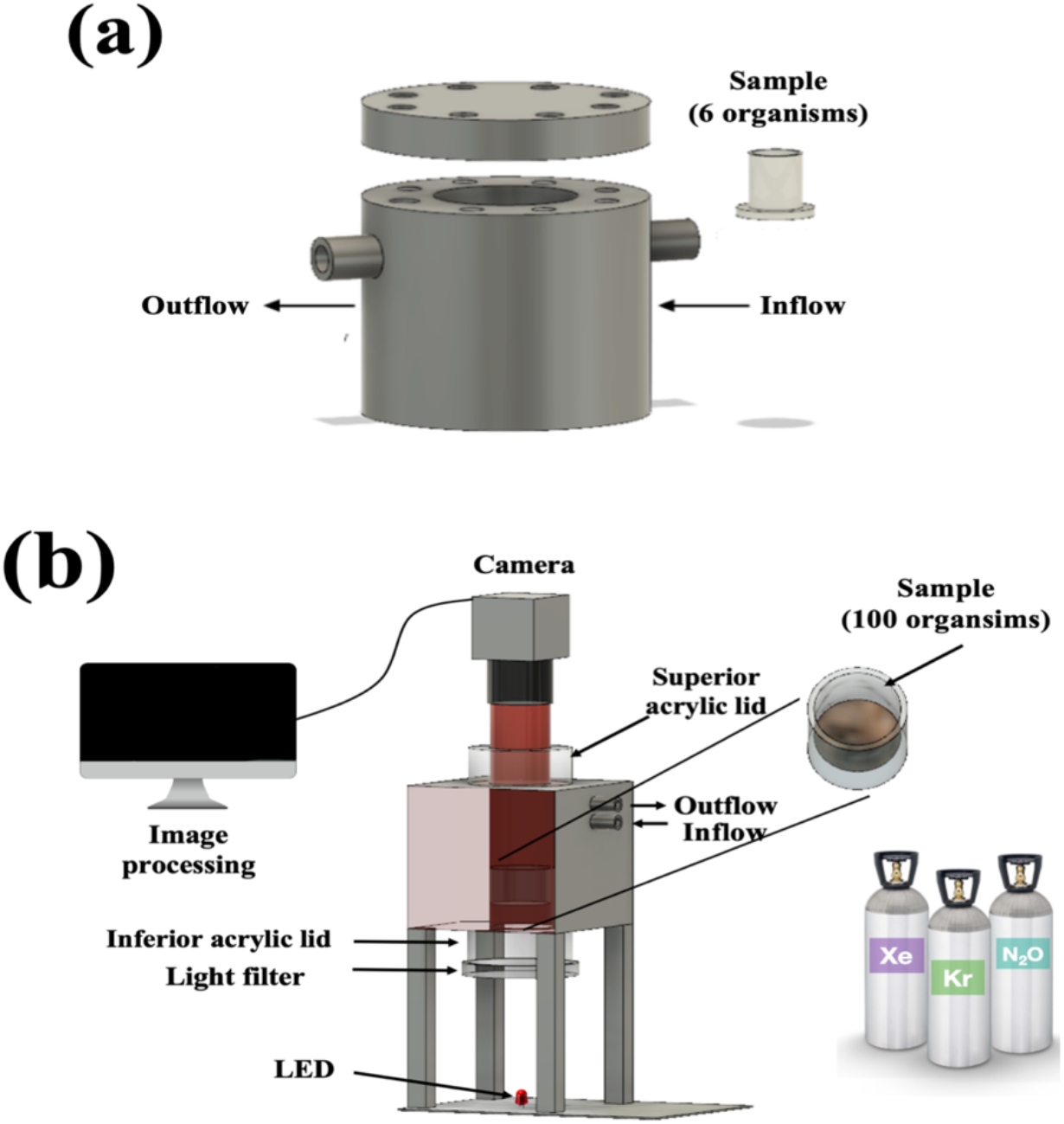
Experimental setup. (a) stainless-steel chamber designed to support high pressures. (b) Experimental setup of anesthesia assay. A red diode led and a light diffusor provide a harmful stimulus for daphnids. The thick acrylic lids allow image recording inside the high pressure chamber. The sample contains around 100 daphnids in free swimming in 10 ml of medium.

### 2.5 Anesthesia assay

The capacity of xenon, krypton and nitrous oxide to suppress motor response after a harmful stimulus was evaluated in DM. For this assay a set of one hundred daphnids 2-4 weeks old were placed into a glass beaker with 10 ml of reconstituted water.^22^ The beaker was placed in an aluminum chamber, see Figure 1b. The chamber has two thick acrylic lids to permit a correct illumination and observation. A diode led was placed at the bottom of the chamber to provide a strong light, which acted as a noxious stimulus.^21^ To evaluate changes in the collective swimming, through the top lid films were captured using a PixeLINK^®^ camera, model PL-B742U (sensor ON Semi MT9M00, USA), operated at 4 frames per second at a 1024×768 pixels resolution. Due to the hydrophobic characteristics of xenon and krypton it was necessary to introduce them in the medium by pressure. The pressures tested were: for xenon 0-14; for krypton 0-44; and for nitrous oxide 0-16 atmospheres. The time of record for each pressure was 30 s, which was an adequate time to observe changes in the collective swimming, see an example in the Supplementary Material. Once the films are saved, the pressure was rapidly increased to the next value.

### 2.6 Image processing

The captured films were transferred to a computer to process them and create separated images. These were analyzed using the ImageJ software 1.46 with an autocorrelation plugin to determine the daphnids motility response (DMR) in the center of successive frames (*t_n_*), respect to the first frame taken at t_0_ (see Figure 2). The model we used to fit the autocorrelation data is a modification of a previously model employed by us in sperm swimming ^26,27^. The values for the autocorrelation C_I_ go from 0 to 1: 0 corresponds to a high motility response after a noxious stimulus while 1 to low motility (anesthetized animals). The autocorrelation values, for different pressures, were adjusted to stretch exponentials to obtain the motility parameter (τ). It is based on the following expression:

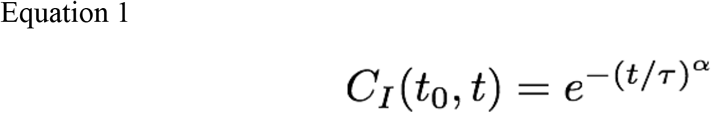

where α is a value that goes from 0.05 to 0.3. Subsequently, a new parameter was employed to characterize the motor response of the organism in presence of a harmful stimulus.

**Figure 2.**
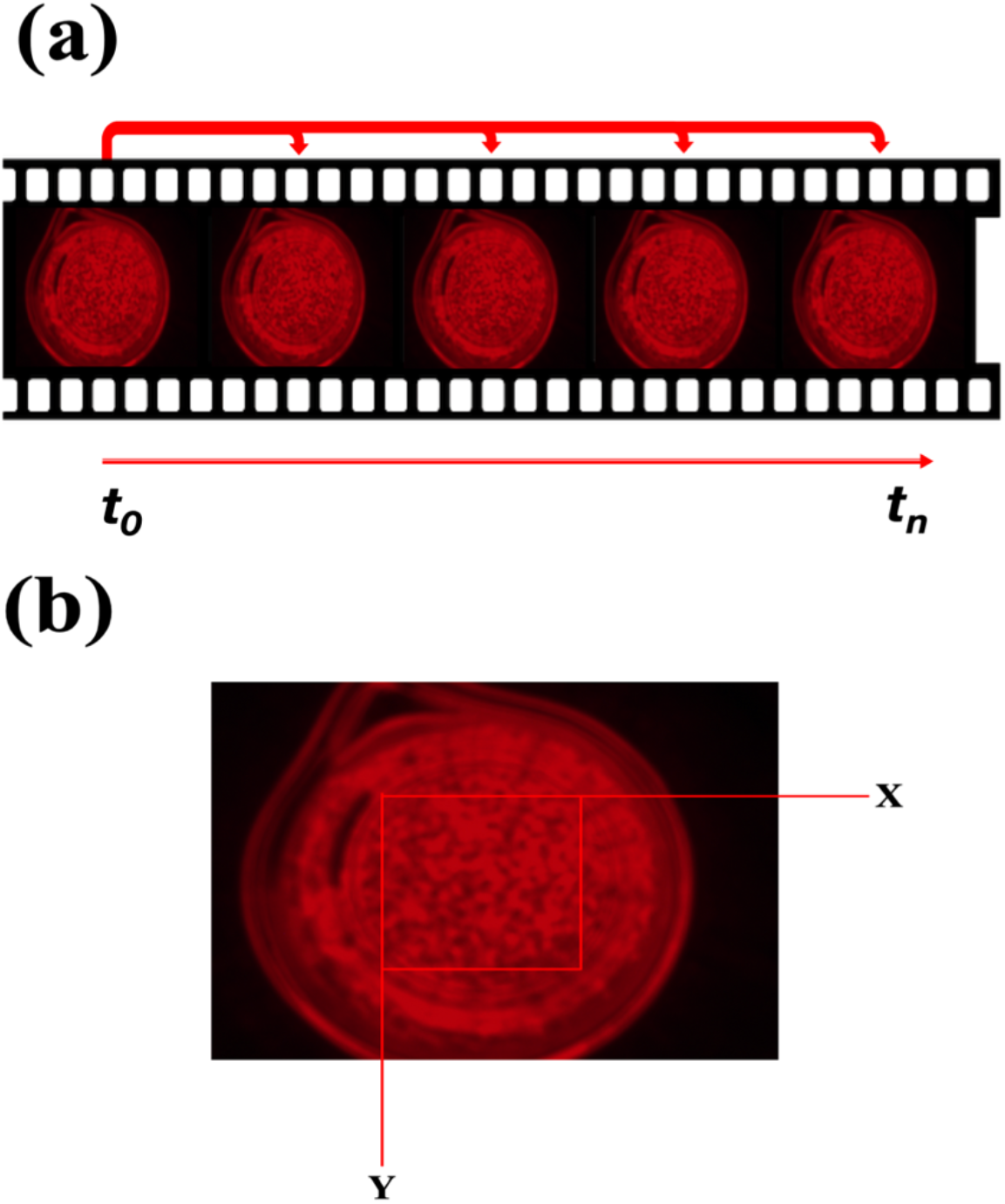
Image processing (a). The first image is taken a t_0_ and subsequent frames at t_n_. (b) Selected region of 250 x 250 pixels at the central part of the frames. The intensity matrix is obtained with those pixels and used to obtain the autocorrelation values.

The daphnids motility response DMR is given by the following equation:

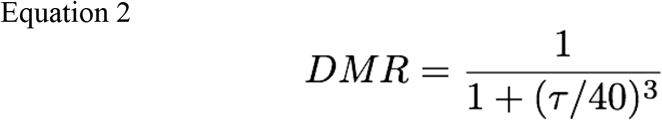

which is bounded between 0 and 1: 1 represents maximum motility and 0 lack of motility. It is important to remark that the values of 40 and 3 in the denominator were obtained, by trial an error, to map low values of τ to 1 and high to 0, see section 3.2.

## 3 Results

### 3.1 Hyperbaric tolerance assay

First, the tolerance test showed that DM is a viable model to be subjected at hyperbaric conditions. Indeed, our measurements showed a high survival of the daphnids to an elevated nitrogen pressure: 95% at 40 atm and 88% at 90 atm. The assay was realized during 1 h. Is important to mention that the subsequent depressurization of the chamber provoked the formation of macroscopic bubbles in the DM’s body. The bubbles subsequently dissolved in the medium and did not compromise their tissues.

### 3.2 Anesthesia assay

As previously mentioned, the anesthetic properties of krypton, xenon, and nitrous oxide were evaluated on *D. magna*. We used the autocorrelation of images to compare changes in the motility. An autocorrelation value of 0 means full movement, which represents a normal response. When this parameter is close to 1, the motility is lost. Krypton presents the lowest capacity to induce immobility, even at high pressures. We subjected the daphnids to pressures from 1 to 44 atm, see figure 3a. The anesthetic effect produced by nitrous oxide is shown in figure 3b. Note that those experiments were carried out at lower pressures 1-10 atm. Finally, the noble gas xenon was supplied at pressures from 1 to 14 atm, see figure 3c.

**Figure 3.**
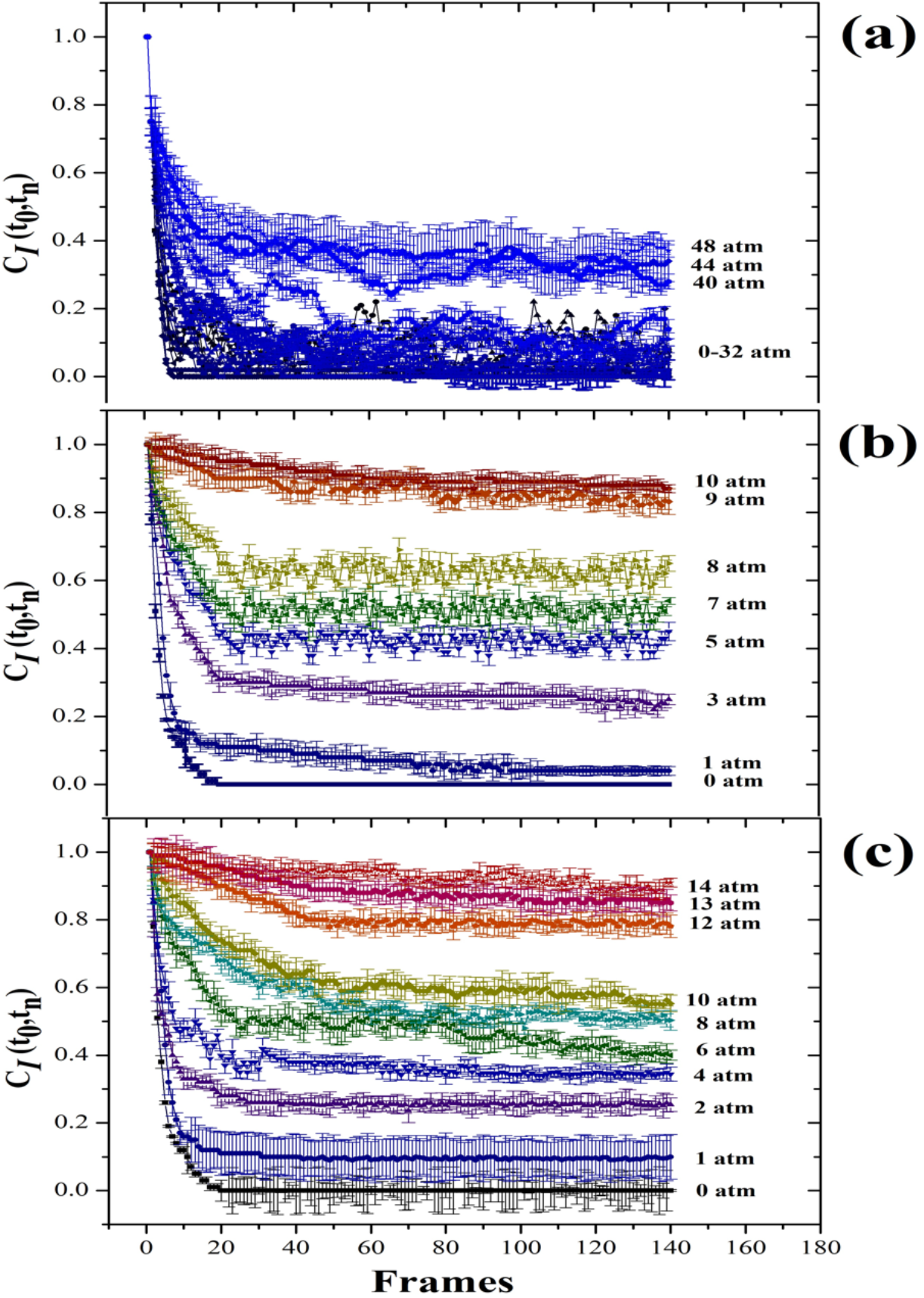
Autocorrelation values as a function of captured frame. Note that the values go from 1 to 0. A sudden drop from 1 to 0 (see for instance the experiments at 0 atm) means that the animals swim normally. As the pressure increases, the autocorrelation values reduce less and less, indicating that the daphnids get immobilized more and more. 1 represents a lack of response to the light harmful stimulus. (a) After exposure to krypton the diminution of movement slightly reduced on *D. magna*, even at pressures near 48 atm. (b) Nitrous oxide showed higher capacity to suppress motility at pressures greater than 3 atm, reaching total immobility at 10 atm. (c) Xenon suppresses motility in *D. magna* at pressures higher than 6 atm. The total immobility was reached at 11 atm.

The data captured and shown in figure 3 give an immediate appraisement of the gases effects on DM. However, as discussed before, two useful parameters are defined to better assess the observed changes. The first parameter τ, see equation 1, allows us to observe the velocity decay of each autocorrelation curve. Small values (1-100) mean large motility. Large values (>100) represent a considerable diminution on the motility. In figure 4, we plot the values of τ as a function of pressure for the tree cases. In a first glance, we can observe the difference between krypton and the other gases.

**Figure 4.**
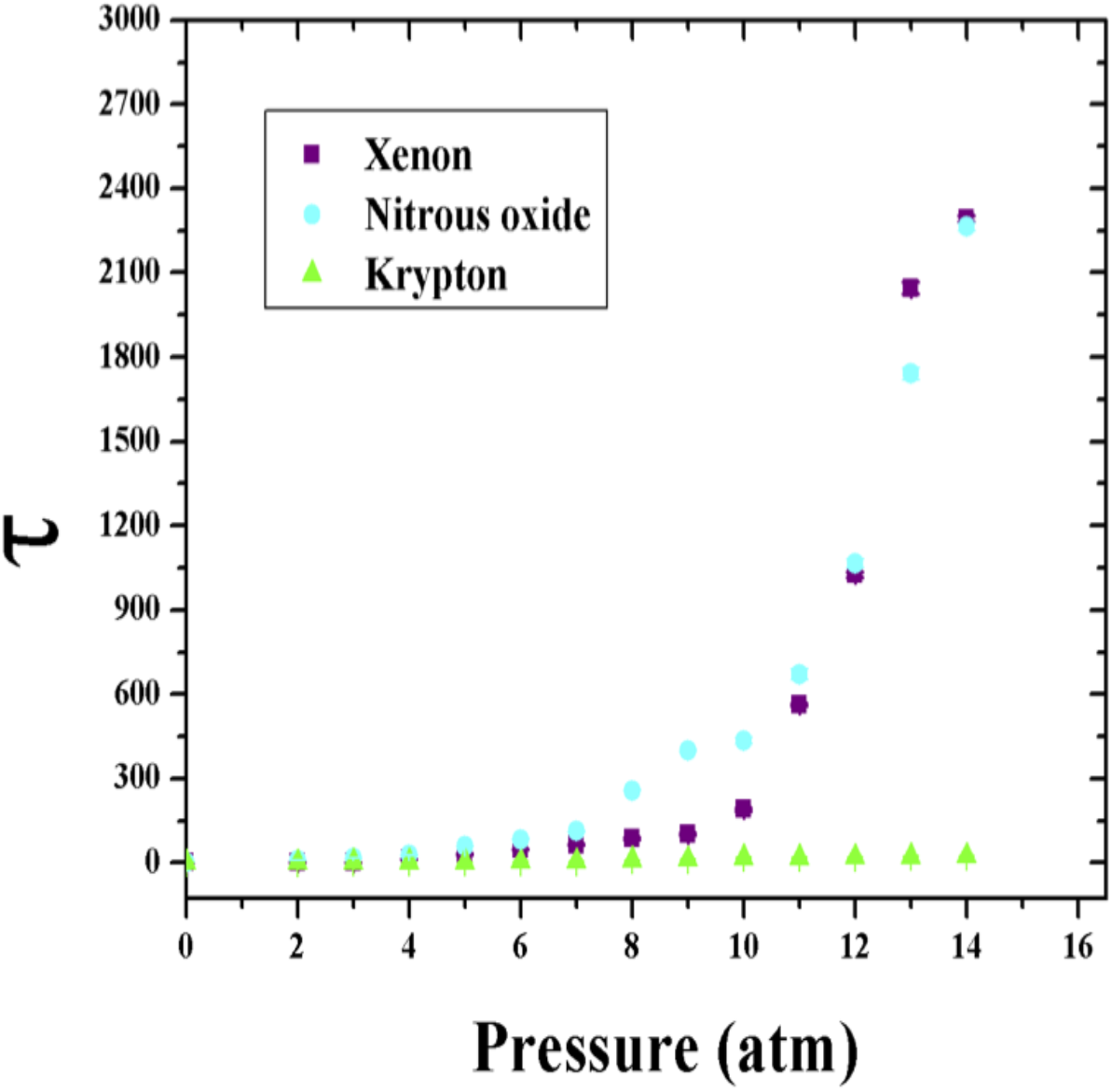
The parameter τ vs pressure for the three gases. Krypton clearly shows a small effect in the motility for all pressures used in the experiment. Xenon and nitrous oxide show an increment in the parameter τ, indicting a change from fast (high motility) to low (small motility) decay in the exponential decaying curves of Fig. 3. The error bars correspond to three different experiments.

According to equation 2, the next step was to determine the change of motility as a function of pressure in order to obtain a dose-response curve. This parameter, DMR, is plotted as a function of pressure, giving a much better appraisal of the anesthesia effects, see figure 5. For example, we found that the EP50 was 5.88 for xenon and for nitrous oxide 4.78 atm.

**Figure 5.**
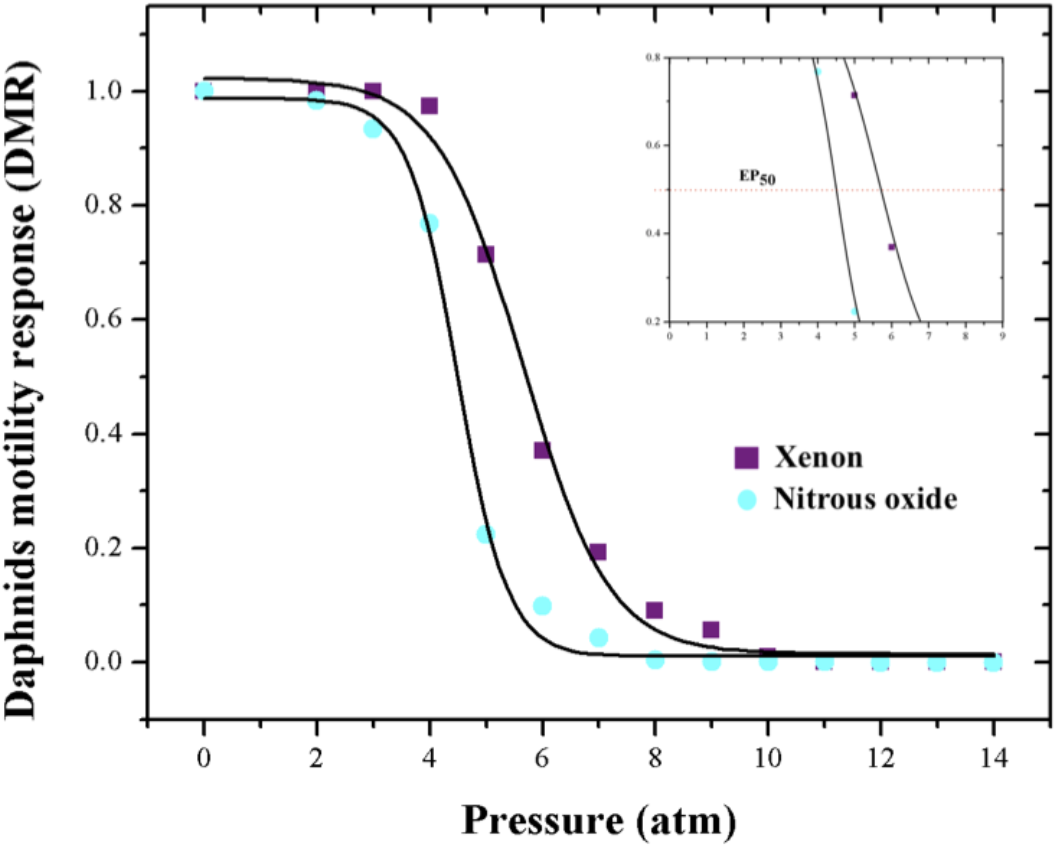
DMR vs pressure for xenon and nitrous oxide. Both reduce the response of DM to a harmful stimulus. Nitrous oxide reduces the 50 % of movement in the invertebrates at 4.78 atm. The total immobility induced by this gas was reached at around 10 atm. Xenon inhibits the motor response in the 50% of the organisms at 5.88 atm. Likewise, this gas fully suppresses the response at around 11 atm.

## 4 Discussion

We carried out experiments using a novel system to test the anesthetic potency of three gases: krypton, nitrous oxide and xenon, on the invertebrates *Daphnia magna*. We developed a simple technique to study in real time, and under pressure, sufficiently large populations of the organisms by using an autocorrelation method.^26,27^ This technique allowed us to assess with high confidence the motor response of the organism under hyperbaric conditions, see movie in the Supplementary Material. We show that the aquatic animal stands as a good candidate to test the anesthetic properties of gases with no physiological risk. Gases dissolve in the aqueous medium by Henry’s law, which states that the amount of dissolved gas in a liquid is proportional to its partial pressure above it.^28^ Our device permits us to dissolve the three gases in the reconstituted water and expose them to the animals. We found that once nitrous oxide and xenon are forced to enter and dissolve in the medium, they produce significative effects in the motor response. Krypton shows the lowest anesthetic effect in *D. magna*, even at a high pressure. Our results are in line we previous results.^29^ We discarded an immobilizing effect produced by pressure by carrying out a control using nitrogen (which is a gas with no anesthetic properties). Moreover, the fact krypton does not produce any loss of motility at pressures around 15 atm (the anesthetic pressures for nitrous oxide and xenon), reinforces the control experiment done with nitrogen. The mechanism of *D. magna* to uptake gases in its aqueous medium remains uncertain. Nevertheless, the processes for blood oxygenation may explain how such gases exchange in the blood flow reaching the nervous system to induce immobility. Oxygen exchanges in *D. magna* occurs near the posterior margin of carapace and in the part of the head.^30^ Alterations produced by low levels of oxygen were discarded, due to the faculty of DM to adapt to a wide range of low-oxygen conditions^31,32^ and also based on the control experiment using high pressures of nitrogen.

The crustacean *Daphnia magna* presents a complex nervous system and not all the receptor have been reported while the use of clones reduced the influence of genetic variations. We presumed that nitrous oxide and xenon would produce motor effects that resembled those observed using volatile anesthetics.^21^ The effects of nitrous oxide and xenon suggest the inhibition of the N-methyl-D-aspartate receptor (NMDAR) in *D. magna*. However, the mild effect seen with krypton cannot be linked to this action mechanism. The standard method to evaluate the potency of a general anesthetic in mammals is the minimal alveolar concentration (MAC), which represents the quantity of an inhalable drug that must exist in the lungs to suppress the motor response after a harmful stimulus in the 50% of a population. It is known that this state is reached once the drug acts on the spinal cord of the animal, been the primary site mediating immobilization.^33^ This effect was described previously after probing that the precollicular decerebration and the complete section of the upper thoracic spinal cord reduce the capacity of sevoflurane to suppress movement in rats. ^34,35^

In summary, we report experiments where three gases (xenon, nitrous oxide, and krypton) are dissolved by exerting pressure on an aquatic medium with daphnid organisms. Nitrous oxide and xenon showed total inhibition of the motor response to harmful stimulus, which could be explained by the inhibition of the N-methyl-D-aspartate receptor (NMDARs).^36–38^ Anesthesia was reversible in short times as soon the pressure is released. The results for these two gases fall in a different range of MACs established for humans.^39,40^

In the context of an increasing number of studies reporting behavioural effects in non-murine animals induced by drugs ^41–43^, we hope the present work may contribute to broaden the advantages to use the motility of aquatic animal models.

## 5 Conclusion

In this study we employed the aquatic organism *D. magna* as an animal pain model for evaluating the anesthetic effects of gases. This crustacean presents the advantage of having a high reproduction rate with low genetic variability. Furthermore, the experimental setup used in the experiments allows us to make a very good statistics and opens the possibility for assays in other aquatic animal models to study suppression of painful sensations. Despite the absence of a defined spinal cord, we probe that this invertebrate organism responds effectively to general anesthetics as vertebrate animals do, suggesting a different site of action in the inhibition of motor responsiveness.

## Author contribution

Study design: K.C.M.R.S and J.C.R.S.

Carried out the experiments: K.C.M.R.S and J.C.R.S.

Data acquisition: K.C.M.R.S, J.C.R.S.

Data analysis: K.C.M.R.S

Drafting of the manuscript: K.C.M.R.S and J.C.R.S.

## Founding

This work has been supported by grants FC-1132 (Conacyt) and FIDSC-100, Mexico.

K.C.M.R.S. acknowledges a scholarship by Conacyt, Mexico.

---

**Table 1.** Comparative of the three gases at different pressures. The parameter τ gives the velocity decay of the autocorrelation curves. The parameter DMR gives the effect on the suppression of motility in the organisms.

| Xe<br>(atm) | $\tau$ | $\Delta\tau$ | DMR | N<sub>2</sub>O<br>(atm) | $\tau$ | $\Delta\tau$ | DMR | Kr<br>(atm) | $\tau$ | $\Delta\tau$ | DMR |
| --- | --- | --- | --- | --- | --- | --- | --- | --- | --- | --- | --- |
| 0 | 1.2 | 0.43 | 0.999 | 0 | 2.1 | 0.02 | 0.999 | 0 | 1.0 | 0.01 | 0.999 |
| 2 | 1.4 | 0.30 | 0.999 | 2 | 10.3 | 0.01 | 0.983 | 2 | 1.6 | 0.01 | 0.999 |
| 3 | 1.7 | 0.02 | 0.999 | 3 | 16.5 | 0.02 | 0.934 | 4 | 2.5 | 0.29 | 0.999 |
| 4 | 12.1 | 0.02 | 0.973 | 4 | 26.8 | 1.00 | 0.768 | 8 | 2.9 | 0.23 | 0.999 |
| 5 | 29.5 | 1.32 | 0.713 | 5 | 60.6 | 0.84 | 0.223 | 12 | 3.4 | 0.13 | 0.999 |
| 6 | 47.8 | 1.63 | 0.369 | 6 | 83.7 | 1.53 | 0.098 | 16 | 3.6 | 0.88 | 0.99 |
| 7 | 64.6 | 2.34 | 0.191 | 7 | 113.1 | 2.43 | 0.042 | 20 | 5.4 | 1.00 | 0.99 |
| 8 | 86.5 | 3.21 | 0.09 | 8 | 257.6 | 2.21 | 0.003 | 24 | 10.2 | 1.00 | 0.98 |
| 9 | 102.4 | 5.21 | 0.05 | 9 | 399.9 | 5.53 | 0.001 | 28 | 12.3 | 2.01 | 0.97 |
| 10 | 190.2 | 5.12 | 0.009 | 10 | 434.4 | 12.99 | 0.001 | 32 | 18.4 | 1.85 | 0.91 |
| 11 | 562.2 | 4.51 | 0.003 | 11 | 669.3 | 19.05 | 0.000 | 36 | 19.0 | 2.13 | 0.90 |
| 12 | 1025 | 42.92 | 5.9427E-05 | 12 | 1065.3 | 15.13 | 5.2932E-05 | 40 | 23.4 | 1.43 | 0.89 |
| 13 | 2291.3 | 22.12 | 5.32E-06 | 13 | 1740.2 | 22.43 | 1.2144E-05 | 44 | 26.1 | 1.45 | 0.85 |
| 14 | 2289.9 | 12.12 | 5.33E-06 | 14 | 2263 | 13.13 | 5.522E-06 | 48 | -- | -- | -- |

## References

1. Aeschbach-Hertig W, Peeters F, Beyerle U, Kipfer R. Interpretation of dissolved atmospheric noble gases in natural waters. Water Resources Research 1999; 35: 2779–2792.

2. Hobbhahn J, Wiesner G, Taeger K. Occupational exposure and environmental pollution: the role of inhalation anesthetics with special consideration of sevoflurane. Der Anaesthesist 1998; 47: 77–86.

3. Vollmer MK, Rhee TS, Rigby M, Hofstetter D, Hill M, Schoenenberger F, Reimann S. Modern inhalation anesthetics: Potent greenhouse gases in the global atmosphere. Geophysical Research Letters 2015; 45: 1606–1611.

4. Marier JR. Halogenated hydrocarbon environmental pollution: The special case of halogenated anesthetics. Environmental research 1982; 28: 212–239.

5. Pothmann W, Shimada K, Goerig M, Fuhlrott M. Pollution of the workplace by anesthetic gases. Causes and prevention. Der Anaesthesist 1991; 40: 339–346.

6. Weber NC, Frädorf J, Ratajczak C, Grueber Y, Schlack W, Hollmann MW, Preckel B. Xenon induces late cardiac preconditioning in vivo: a role for cyclooxygenase 2?. Anesthesia & Analgesia 2008; 107: 1807–1813.

7. Hartlage MAG, Berendes E, Van Aken H, Fobker M, Theisen, M, Weber TP. Xenon improves recovery from myocardial stunning in chronically instrumented dogs. Anesthesia & Analgesia 2004; 99: 655–664.

8. Preckel B, Müllenheim J, Moloschavij A, Thämer V, Schlack W. Xenon administration during early reperfusion reduces infarct size after regional ischemia in the rabbit heart in vivo. Anesthesia & Analgesia 2000; 91: 1327–1332.

9. Olivot JM, Mlynash M, Zaharchuk G, Strak M, Bammer R, Schwartz N, Albers GW. Perfusion MRI (Tmax and MTT) correlation with xenon CT cerebral blood flow in stroke patients. Neurology 2009; 72: 1140–1145.

10. Wintermark M, Thira JP, Maeder P, Schnyder P, Meuli R. Simultaneous measurement of regional cerebral blood flow by perfusion CT and stable xenon CT: a validation study. American journal of neuroradiology 2001; 22: 905–914.

11. Schoenborn BP. Binding of xenon to horse haemoglobin. Nature 1965; 208: 760–762.

12. Hecker K, Baumert JH, Horn N, Rossaint R. Xenon, a modern anaesthesia gas. Minerva anestesiologica 2004; 70: 255–260.

13. Urban BW. The site of anesthetic action. In Modern Anesthetics 2008; 127: 3–29.

14. Lynch C, Baum J, Tenbrinck R, Weiskopf RB. Xenon anesthesia. The Journal of the American Society of Anesthesiologists 2000; 92: 865–870.

15. Goto T, Saito H, Shinkai M, Nakata Y, Ichinose F, Morita S. Xenon provides faster emergence from anesthesia than does nitrous oxide-sevoflurane or nitrous oxide-isoflurane. Anesthesiology 1997; 86: 1273–1278.

16. Jordan BD, Wright EL. Xenon as an anesthetic agent. AANA J 2010; 78: 387–392.

17. Khan KS, Hayes I Buggy DJ. Pharmacology of anaesthetic agents II: inhalation anaesthetic agents. Continuing Education in Anaesthesia, Critical Care & Pain 2014; 14: 106–111.

18. Gall O, Annequin D, Benoit G, Van Glabeke E, Vrancea F, Murat I. Adverse events of premixed nitrous oxide and oxygen for procedural sedation in children. The Lancet 2001; 358:1514–1515.

19. Utsumi J, Adachi T, Miyazaki Y, Kurata J, Shibata M, Murakawa M, Mori K. The effect of xenon on spinal dorsal horn neurons: a comparison with nitrous oxide. Anesthesia & Analgesia 1997; 84: 1372–1376.

20. Derwall M, Coburn M, Rex S, Hein M, Rossaint R, Fries M. Xenon: recent developments and future perspectives. Minerva anestesiologica 2009; 75: 37–45.

21. Mckenzie, JD, Calow P, Nimmo WS. A model to test the potency of inhalation anaesthetics. BJA: British Journal of Anaesthesia 1989; 63: 489–491.

22. Robledo-Sánchez KCM, Guevara-Pantoja FJ, Ruiz-Suárez JC. Video-tracking and high-speed bright field microscopy allow the determination of swimming and cardiac effects of D. magna exposed to local anaesthetics. Science of the Total Environment 2019; 691: 278–283.

23. Rivetti C, Campos B, Barata C. Low environmental levels of neuro-active pharmaceuticals alter phototactic behaviour and reproduction in Daphnia magna. Aquatic Toxicology 2016; 170: 289–296.

24. Barata C, Markich SJ, Baird DJ, Soares AM. The relative importance of water and food as cadmium sources to Daphnia magna Straus. Aquatic Toxicology 2002; 61: 143–154.

25. Kluttgen B, Ratte HT. Effects of different food doses on cadmium toxicity to Daphnia magna. Environmental Toxicology and Chemistry: An International Journal 1994; 13: 1619–1627.

26. Sierra-Valdez FJ, Cisneros-Mejorado AJ, Sánchez Herrera, Ruiz-Suárez JC. A thermal study of cellular motility by optical time-resolved correlation. Review of Scientific Instruments. 2012; 83: 044302.

27. Ríos-Ramírez M, Espinoza JH, Ruiz-Suárez JC, Mercado-Uribe H. The effect of green light on the motility of mouse sperm at two different temperatures. Photochemical & Photobiological Sciences. 2019; 18: 2893–2900.

28. Kennan RP, Pollack GL. Pressure dependence of the solubility of nitrogen, argon, krypton, and xenon in water. The Journal of chemical physics 1990; 93: 2724–2735.

29. Cullen SC, Gross EG. The anesthetic properties of xenon in animals and human beings, with additional observations on krypton. Science 1951; 113: 580–582.

30. Pirow R, Wollinger F, Paul RJ. The sites of respiratory gas exchange in the planktonic crustacean Daphnia magna: an in vivo study employing blood haemoglobin as an internal oxygen probe. Journal of Experimental Biology 1999; 202: 3089–3099.

31. Weider LJ, Lampert W. Differential response of Daphnia genotypes to oxygen stress: respiration rates, hemoglobin content and low-oxygen tolerance. Oecologia 1985; 65: 487–491.

32. Rüdiger JP, Colmorgen M, Pirow R, Chen YH, Tsai MC. Systemic and metabolic responses in Daphnia magna to anoxia. Comparative Biochemistry and Physiology Part A: Molecular & Integrative Physiology 1998; 120: 519–530.

33. Sonner JM, Antognini JF, Dutton RC, Flood P, Gray AT, Harris RA, Trudell, J. Inhaled anesthetics and immobility: mechanisms, mysteries, and minimum alveolar anesthetic concentration. Anesthesia & Analgesia 2003; 97: 718–740.

34. Rampil IJ, Mason P, Singh H. Anesthetic potency (MAC) is independent of forebrain structures in the rat. Anesthesiology 1993; 78: 707–712.

35. Rampil IJ. Anesthetic potency is not altered after hypothermic spinal cord transection in rats. Anesthesiology 1994; 80: 606–610.

36. Nagele P, Metz LB, Crowder CM. Nitrous oxide (N2O) requires the N-methyl-D-aspartate receptor for its action in Caenorhabditis elegans. Proceedings of the National Academy of Sciences 2004; 101: 8791–8796.

37. Franks NP, Dickinson R, De Sousa SLM, Hall NA, Lieb WR. How does xenon produce anaesthesia? Nature 1998; 396: 324–324.

38. Goto T, Nakata Y, Saito H, Ishiguro Y, Niimi Y, Suwa K, Morita S. Bispectral analysis of the electroencephalogram does not predict responsiveness to verbal command in patients emerging from xenon anaesthesia. British journal of anaesthesia 2000; 85: 359–363.

39. Jevtović-Todorović V, Todorovć SM, Mennerick S, Powell S, Dikranian K, Benshoff N, Olney JW. Nitrous oxide (laughing gas) is an NMDA antagonist, neuroprotectant and neurotoxin. Nature medicine 1998; 4: 460–463.

40. Hornbein TF, Winter PM, Smith G, Wetstone D, Smit KH. The minimum alveolar concentration of nitrous oxide in man. Anesthesia and Analgesia 1982; 61: 553–556.

41. Imeh-Nathaniel A. et a.. Effects of chronic cocaine, morphine and methamphetamine on the mobility, immobility and stereotyped behaviors in crayfish. Behavioural Brain Research 2017; 332: 120–125.

42. Nathaniel TI, Panksepp J, Huber R. Drug-seeking behavior in an invertebrate system: Evidence of morphine-induced reward, extinction and reinstatement in crayfish. Behavioural Brain Research 2009; 197: 331–338.

43. Lieschke GJ and Currie PD. Animal models of human disease: zebrafish swim into view. Nature Review Genetics 2007; 8: 353–367.

